# CRISPR-based multi-locus real-time tracking reveals single chromosome dynamics and compaction

**DOI:** 10.1101/2022.02.01.478681

**Authors:** Yu-Chieh Chung, Madhoolika Bisht, Li-Chun Tu

**Affiliations:** Department of Biological Chemistry and Pharmacology, The Ohio State University, Columbus, OH, USA; Department of Molecular Genetics, The Ohio State University, Columbus, OH, USA; Center for RNA Biology, The Ohio State University, Columbus, OH, USA; The Ohio State University Comprehensive Cancer Center, The Ohio State University, Columbus, OH, USA

**Keywords:** Chromatin dynamics, Single molecule microscopy, CRISPR, Live-cell imaging

## Abstract

In eukaryotic nuclei, individual chromosomes occupy discrete three-dimensional spaces with little overlap. Dynamic chromatin organization instantly influences DNA accessibility through modulating local macromolecular density and interactions, driving changes in transcription activities. Chromatin anchoring to nuclear landmarks often leads to massive reorganization and motion changes. Chromatin dynamics has been reported to be locally confined but contributes to the large-scale coherent chromatin motion across the entire nucleus. However, the dynamics and compaction of chromosomal sub-regions along a single chromosome are not well-understood. In this study, we combined quantitative real-time single-molecule fluorescence microscopy, CRISPR-based genomic labeling, biophysical analysis and polymer models to characterize the dynamics of specific genomic loci and chromatin-nuclear landmark interactions on human chromosome 19 in living cells. Precise genomic labeling allows us to dissect loci motions and chromatin elasticity on a single chromosome basis. We found that the dynamics of genomic loci were all subdiffusive but varied at different regions along the chromosome. The mobility of genomic loci was similar among interior chromosomal loci but deviated for the loci at pericentromeric and near-telomeric regions. Tighter compaction on chromosome 19 long arm, compared to the short arm was observed, which may correlate to more active genes on the short arm than the long arm, shown by our RNA-seq analysis. The strong tethering interaction was found for loci at the pericentromeric region, suggesting a higher degree of local condensation, perhaps through stronger interactions or association between pericentromeric regions and their microenvironments, such as chromatin-nuclear body association through sequence-specific domains on DNA.

## Introduction

Chromatin compaction, localization, and dynamics are orchestrated for precise cellular processes. Mis-regulation of chromatin organization has been shown to associate with diseases including developmental defects and cancers(1, 2). Interphase chromatin is non-randomly and hierarchically organized in the cell nucleus. The chromatin compaction affects transcription control. Transcriptionally active chromatin is generally less packed than transcriptionally inactive chromatin due to the chromatin remodeler activities at transcribed gene loci (3). The gene density and number of active genes on a chromosome differs based on the cell type and its diseased state. Our RNAseq analysis showed that the number of active genes is less in the q arm compared to the number of active genes in the p arm on human chromosome 19. How compaction levels differ in the p and q arm of a chromosome remains unexplored.

Dynamic nuclear compartments affect the biophysical properties of the regional nuclear macromolecules (4). The peripheral nucleosomes showed lower mobility compared to the interior nucleosomes in the nucleus (5). Our previous work showed that genomic loci at telomere have higher mobility compared to the locus mobility in interior chromosome and pericentromeric regions (6). The different dynamics between genomic loci are expected when distinct nuclear microenvironments are present or the tethering of chromatin to nuclear landmarks occurs. For example, the centromeric region of chromatin is primarily more condensed compared to condensation levels at other regions of a chromosome (7). Chromatin domains in pericentromeric regions tends to cluster together and have a higher level of methylated histone marks when compare to interior chromosomal regions (8). On the other hand, telomeric regions are bound by highly dynamic telomerase and its cofactors (9). A large number of lysines on histones are hypoacetylated near telomeric regions (10). It raises questions if chromatin mobility and compaction are chromosome- and locus-dependent.

Chromatin can associate with nuclear landmarks, such as nuclear bodies and nuclear lamina, through corresponding associated domains (11). These chromatin-landmark interactions are critical to chromatin localization, organization, dynamics and function in the cell nucleus. For example, gene loci in close proximity to the nuclear lamina were localized to the nuclear periphery and often suppressed. However, measurement of the tethering of chromatin domains to their surrounding nuclear landmarks is challenging in living cells. Genomic locus dynamics is subject to chromatin elasticity as well as the interactions with molecules or landmarks in its microenvironment, such as nuclear bodies, nuclear lamina, and soluble molecules in the nucleoplasm. Changes in motion and location of genomic loci were found to occur in response to transcription activation, DNA damage repair, and terminal differentiation (12-15). Investigating local chromatin dynamics can reveal a wealth of information about chromatin elasticity and its interaction with microenvironments, which is less likely to be interrogated by using techniques that work on static chromatin. Live-cell imaging has provided substantial spatiotemporal insights regarding genomic organization at the single-cell level. The biophysical properties of genomic loci can be correlated to transcription activity and be used to map the inhomogeneous distribution of active and inactive genes. Our CRISPR-Sirius imaging technique is developed to track the real-time chromatin dynamics in 3D nuclear space and is a powerful tool for measuring biophysical prosperities of genomic loci on single chromosomes(16).

CRISPR (clustered regularly interspaced short palindromic repeats) and its associated proteins (Cas) have been repurposed to track movement of genomic loci in living cells (6, 16, 17). Type II CRISPR-Cas9 system is one of the most commonly used CRISPR system for genomic reorganization and imaging(18). Cas9 naturally is an endonuclease which generates double strand breaks when forming stable interactions with guide RNA and DNA. Two mutations, D10A and H840A, were introduced to create the nuclease-dead version of Cas9 (dCas9) to remove the endonuclease activity (19). In CRISPR-Sirius, engineered single guide RNAs (sgRNAs) with multiplexed RNA aptamers (PP7 and MS2) were used with its fluorescently-tagged RNA coat proteins to track the movement of dual loci simultaneously on a single chromosome (16). Advantages of CRISPR-based imaging techniques includes: (1) single-chromosome studies available with no modification of endogenous DNA sequences for the labeling; (2) the fluorescence intensity is stable for hours without blinking issues; (3) more than 1000 loci in human genome can be labeled based on the database search results; and (4) higher resolution can be achieved by using the shorter genomic targeting length. For example, two loci that are 4.6 kb apart were resolved by using the labeling region in less than a kilobase length (20).

In this work, we use CRISPR-Sirius to measure the biophysical properties of genomic loci on human chromosome 19 and characterize their variations in different genomic regions along a single chromosome. We measured chromosome compaction by using the scaling exponent of the power-law relationship of the averaged spatial distance and genomic distance of locus pairs. The locus territory, the area covered by locus movement, and the mean squared displacement (MSD) of individual loci were analyzed by the trajectory of the locus movement. To quantify the tethering coupling between chromatin and its local environments, we measured the effective spring constant for individual loci and then extracted the inter-locus and tethering interactions by using the Rouse polymer model and dual-color two-locus tracking data. We established an approach to studying the genomic locus dynamics and its variation along a single chromosome as well as to quantifying the coupling of loci to nearby nuclear landmarks, such as nuclear bodies or lamina, which can serve as a foundation for having a general understanding of chromosome dynamics and nucleus function.

## Results

### Imaging genomic loci on human chromosome 19 by CRISPR-Sirius

Our experimental design for labeling and imaging specific genomic loci is based on our CRISPR-Sirius work (Figure 1A)(16). Two octets of RNA aptamers from bacteriophages, MS2 and PP7, were inserted into separate single guide RNA (sgRNA) scaffolds, to generate sgRNAs used for the simultaneous dual-color labeling of specific genomic loci with high brightness. The aptamers were designed and engineered in three-way junctions for enhancing thermostability which prevents the sgRNA from fast degradation in the human nucleus(16). The MS2 coat protein (MCP) and PP7 coat protein (PCP) were genetically fused to the green fluorescent protein (GFP) and halo tag, respectively, to generate PCP-GFP and MCP-halo tag for the fluorescent tagging of Sirius sgRNAs in live cells. The membrane permeable dye JF549 was used to label the halo tag (21). To probe loci dynamics along chromosome 19, six loci were selected, including a locus near the telomeric region, T2, three loci on the long arm, LA, LH, and LE, and two loci in pericentromeric regions, PR1 and PR2 (Figure 1B). All sgRNAs showed well-targeted pairs in dual-color in U2OS cells (Figure 1C). A foci count of LA showed mostly two and three labeled foci (with an average number of 2.5 foci per cell) were detected in cells when scanning the whole three-dimensional nucleus (Figure S3), which is consistent with the expected numbers of chromosomes in U2OS reported previously (16).

**Figure 1.**
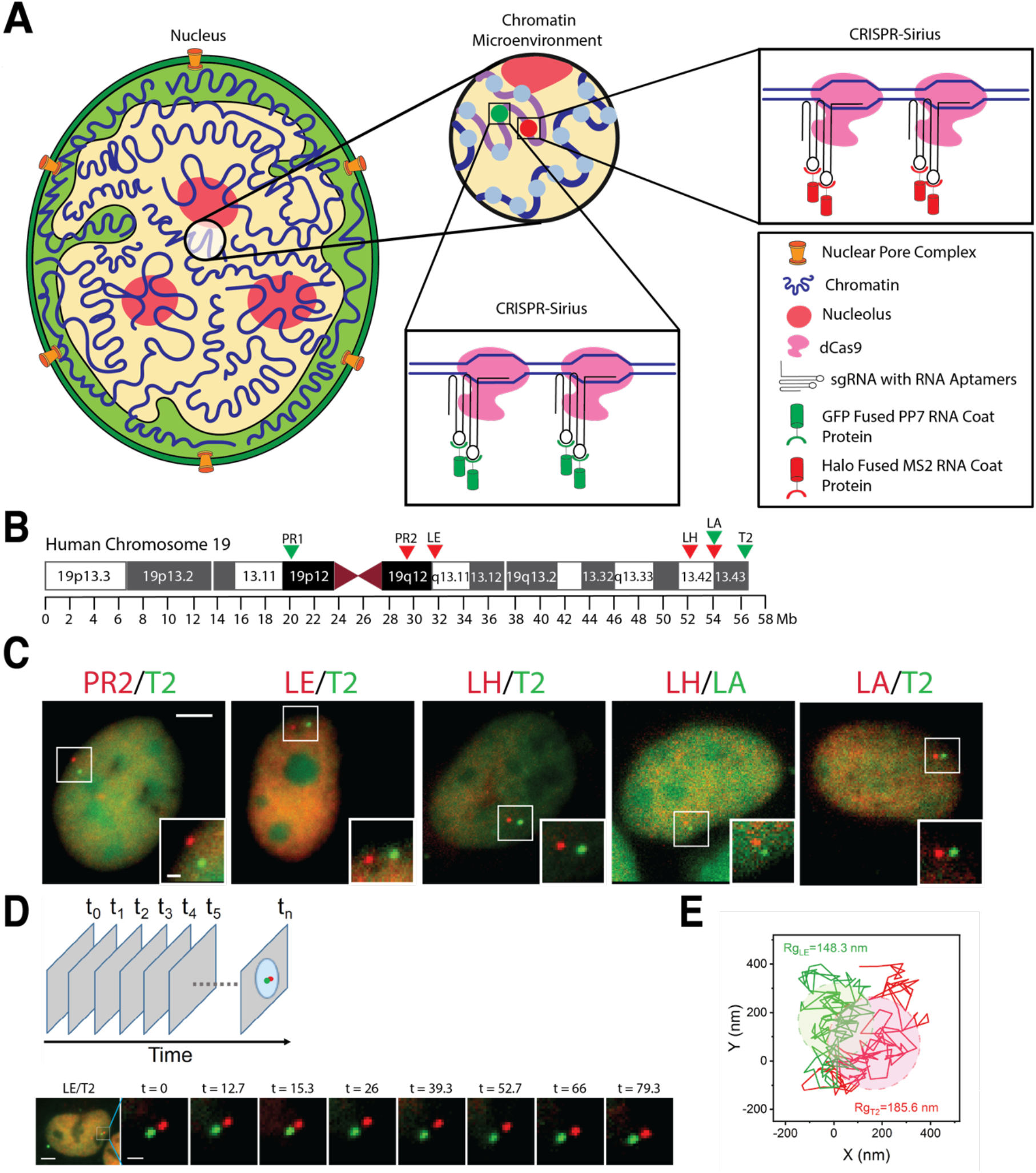
Dual-color imaging and trajectories of locus pairs by CRISPR-Sirius. (A) Diagram of dual-color CRISPR-Sirius system for specific genomic imaging in living cells (16). (B) Genomic location of labeled loci along human chromosome 19. (C) Visualization of dual-color labeled genomic loci in U2OS cells. White boxes indicate locations of labeled loci in the nucleus and are enlarged in the right bottom corners. Scale bar, 5 μm. Bar in the inset, 1 μm. The green and red colors are false colors indicating loci labeled by hU6-sgRNA-Sirius-8XPP7-GFP and mU6-sgRNA-Sirius-8XMS2-halo tag-JF549, respectively. (D) A time-laps experiment of LE-T2 locus pair over 80 seconds (total frame number = 120). Scale bar in the first image, 5 μm. Bar in the enlarged inset, 1 μm. (E) Trajectories shows mobilities of LE and T2 in (D). Gyration radii of trajectories are indicated in similar colors.

### Distinct chromatin compaction levels for chromosome p arm and q arm

Chromatin structure is hierarchical in the 3D nuclear space. Although substantial understanding has been elucidated at the nucleosome scale (~11 nm), large-scale chromatin organization and dynamics in the cell nucleus is still not fully understood due to the complexity. Because the large-scale chromatin organization is influenced by transcription activities and the gene density varying in each chromosome (22), we hypothesize individual chromosomes or chromatin domains on a single chromosome can be organized into different levels of compaction. Our previous study on chromosome 19 p arm showed that the inter-locus distance of locus pairs from kilobases to megabases was positively correlated to the genomic distance but become less or non-correlated when the genomic distance was larger than ~10 Mb (20). What is the compaction level of chromosome 19 q arm? And are the two arms of a single chromosome folded in the same compaction level? These very interesting and important questions remain to be answered.

To measure the compaction of chromosome 19 q arm, we paired genomic loci on the long arm with various genomic distances - 1.93 Mb (LH/A4), 2.69 MB (LA/T2), 4.62 Mb (LH/T2), 25.82 MB (LE/T2), and 29.05 MB (PR2/T2) (Figure 2A). The spatial distance of these locus pairs was measured and plotted against their genomic distance. The compaction level of chromatin is determined by the scaling exponent (δ, compaction exponent) of the power-law relationship between the spatial distance and genomic distance of loci pairs. As shown in Figure 2B, genomic length-dependent compaction was observed again on chromosome 19 q arm. More precisely, δ_4.6Mb_ equals 0.40 at 4.6 Mb genomic distance, δ_25Mb_ equals 0.18 at 25.8 Mb genomic distance, and δ_29Mb_ equals 0.20 at 29 Mb genomic distance (Table S2). Surprisingly, by comparing the compaction exponents at ~25 Mb size, we found that chromosome 19 q arm (δ_25Mb_ = 0.18) (Figure 2B) was packed tighter than the p arm (δ_21Mb_= 0.22)(20). We analyzed the RNA-seq data of U2OS cells and found that although the total number of genes on the q arm (n_q_ = 994) is more than that of genes on the p arm (n_p_ = 763), the number of active genes on the p arm is ~10% more than the active genes on the q arm in U2OS cells (p is 442 vs. q is 400, Figure 2C). Taking the total genes on each arm into account, ~40% of genes on the q arm are active and the active gene percentage on the p arm is 58%. The more active genes indicate a potentially lower nucleosome density and more active transcription factor interactions. These factors were demonstrated to increase chromatin accessibility by DNaseI cleavage and ChIP-seq techniques (3). We reasoned that intense transcription activities lead to a more extended and relaxed chromatin configuration.

**Figure 2.**
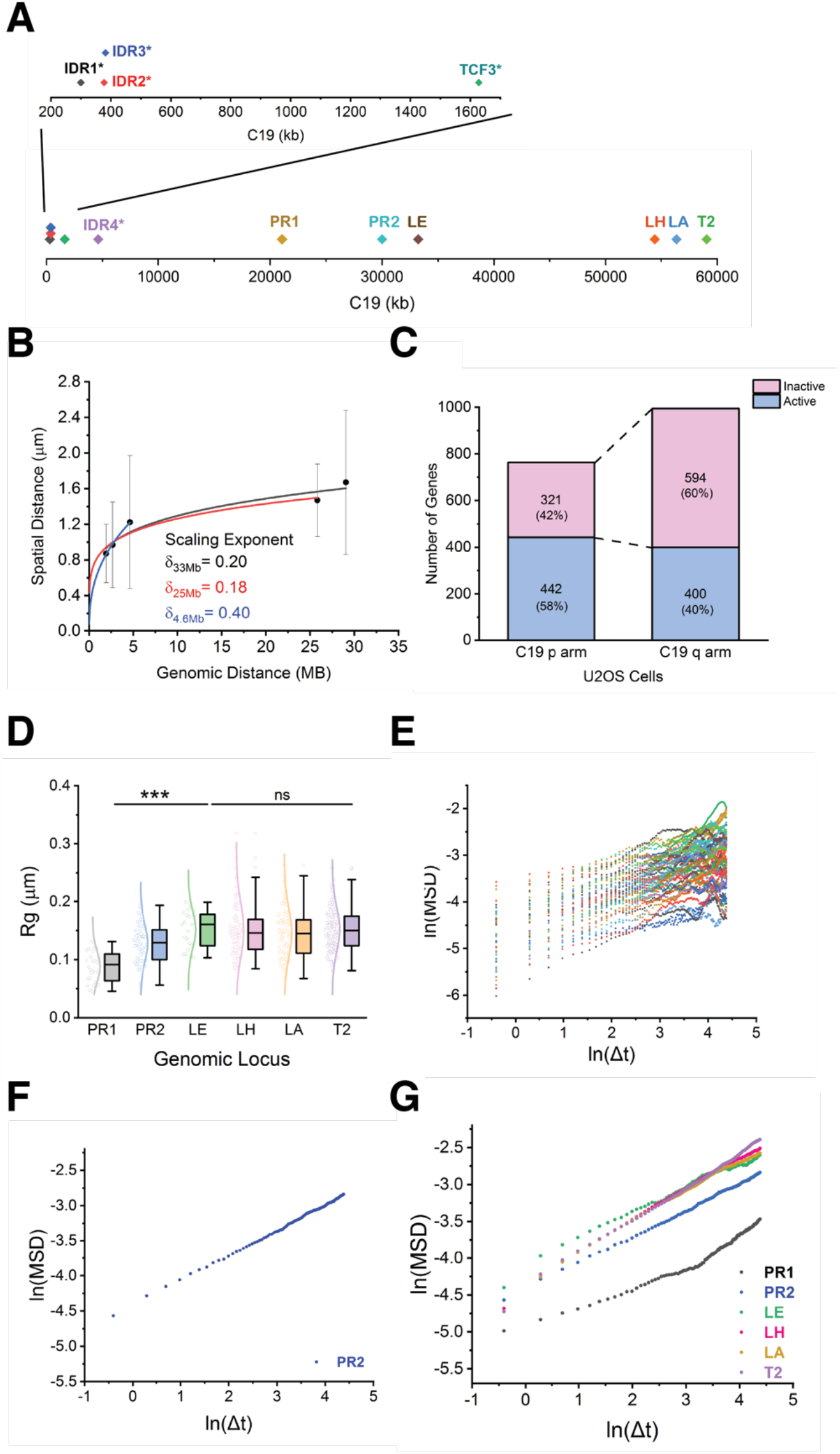
Compaction and dynamics of human chromosome 19. (A) Genomic coordinates of previous reported loci (IDR1, IDR2, IDR3, TCF3, and IDR4, marked with a *) (20) and labeled loci in this study along human chromosome 19. Previous loci only span ~7.8 % of the chromosome and localized on the p arm. (B) Mean spatial distance against genomic distance for chromosome 19 q arm. The scaling exponents δ are given by fitting the data to the power law relationship. The error bars of the plot represent one standard deviation. (C) The RNA-seq analysis of U2OS cells showed the actively transcribed genes verse the inactive genes on the chromosome 19 p and q arms (active genes in blue and inactive genes in pink). (D) Box-and-Whisker plot of trajectory radii of individual loci (PR1, PR2, LE, LH, LA, and T2). Data points and distribution curves are indicated to the left of each box. One-way ANOVA test showed statistically significance (*P* << 0.0005) among the loci in the pericentromeric regions (PR1 and PR2) and the interior locus (LE). No statistical significance (*p* > 0.05) was found among interior loci (LE, LH, LA) and the near-telomeric locus (T2). (E) The time-averaged MSD plot of PR2 locus in a natural logarithm scale (n = 52 trajectories). (F) The essemble-averaged MSD curve from (E). (G) Essemble-averaged MSD curves of all loci, PR1, PR2, LE, LH, LA, T2 (n = 28 trajectories for PR1, 52 trajectories for PR2, 21 trajectories for LE, 77 trajectories for LH, 77 trajectories for LA, and 127 trajectories for T2).

### Characterization of genomic locus dynamics and territory along a single chromosome

We have previously characterized five loci motion – IDR1, IDR2, IDR3, TCF3 and IDR4 - located on human chromosome 19 p arm within genomic coordinate 0 to 4.6 Mb (Figure 2A)(20). The total genomic length of human chromosome 19 is about 59 Mb but our previous study only covered 7.8 % region of the chromosome. By adding six new labeled loci in this work, we expand the coverage of loci over the whole chromosome 19 to improve the understanding of single-chromosome dynamics. The movement of a locus was recorded in consecutive 120 image frames for 80 seconds (Figure 1D) and the trajectory of the locus is given by the time series of its position on each frame (Figure 1E). The lateral localization precision determined by 100 nm coverslip-absorbed Tetraspeck beads was ~ 6 nm in 16 seconds (Figure S1) and ~10 nm in 80 seconds (Figure S2). Localization precisions were slightly larger (~1 nm difference) in the red (561 nm excitation) channel compared to the green channel (488 nm excitation) in 16 seconds but this difference became insignificant when total imaging time is much shorter (4 seconds) (Figure S1A) or longer (80 seconds) (Figure S2). When separating x and y axis, a slightly better (~1 nm difference) localization precision along the y axis compared to the x axis was observed (Figure S1B). The movement of genomic loci can be quantitatively detected and compared by two biophysical parameters – the gyration radius (trajectory radius, R_g_) and mean squared displacement (MSD) of locus trajectory (Figure 2D and 2E, Supplementary Information). The trajectory radius represents the area covered by the trajectory of a genomic locus movement within a given time and can be regarded as “locus territory”. We found similar trajectory radii (Figure 2D) on interior loci with 1.46 x 10^-1^ mm at LA, 1.51 x 10^-1^ mm at LH, and 1.51 x 10^-1^ mm at LE and 1.52 x 10^-1^ mm at the telomeric locus T2. However, the trajectory radii of loci at pericentromeric regions were significantly smaller with 0.88 x 10^-1^ mm at PR1 and 1.28 x 10^-1^ mm at PR2 (Table S1), consistent with the previous trajectory radius result of loci, IDR1, IDR2, IDR3, TCF3 and IDR4 (interior and near-telomeric region on the p arm). This suggests the movement of loci within the pericentromeric regions are more constrained than that of loci within the interior and telomeric regions for both arms of chromosome 19. As a control, we examined the RNA-seq data of human chromosome 19 in U2OS (Materials and Methods). None of the loci used in this research (PR1, PR2, LE, LH, LA and T2) is on an active gene (Table S4). Apart from the locus territory, the mobility of loci can be characterized by MSD. Apparent diffusion constants (D*_app_*) and diffusion exponents (β) were extracted from fitting the data to the power-law MSD function (MSD = 4D*_app_*Dt^β^). The time-averaged MSD of individual trajectories (Figure 2E) and the ensemble-averaged of MSDs of loci over cell population were calculated (Figure 2F and 2G). Consistent with our previous study of genomic loci on chromosome 19 p arm (IDR1, IDR2, IDR3, TCF3, and IDR4) (20), the diffusion exponents were found to be between 0.32 to 0.46 (Table S1), indicating different levels of subdiffusive motions of loci along chromosome 19. The mobility of loci can be seen from their MSD curves (Figure S4). Although MSD curves of T2 near telomeric region overlap with those of interior loci, LA, and LH, initially, T2 was less restricted after 35 seconds, compared to MSDs of LA and LH and showed the highest mobility in a long time period. In a short-time period (less than 10 seconds), the locus LE possesses the highest mobility among others. It is noteworthy that interior and near telomeric loci showed higher mobility than the loci at the pericentromeric region at all time points.

### Chromatin elasticity and tethering to its local environment

We have shown that the dynamics of loci on chromosome 19 is subdiffusive in the previous section and in (20). The subdiffusive dynamics of genomic loci can be modeled as the generalized Langevin equation (23, 24). The genomic locus dynamics is constrained by external forces, mainly including inter-locus interactions, the tethering interaction of a locus to the local microenvironment, such as nuclear landmarks and the frictional force from its surrounding medium. In a short time, the system can be modeled by the normal Langevin equation (25) in which the locus motion can be approximated as a normal diffusion with an effective diffusion constant *D_eff_*, and the local nucleoplasm will be treated as a viscous medium with a friction coefficient g related to effective diffusion constant *D_eff_* by the Einstein relation, g = *K_B_T/D_eff_*. We assume the effective external force applied on the locus obeys Hooke’s law with an effective spring constant *k_eff_* (26). The effective spring constant *k_eff_* can be calculated by the linear regression of step size of locus trajectory versus the relative position of locus to the centroid of the trajectory (25) (Figure 3A and 3B, Supplementary Information). We found that the effective spring constant of the locus at the pericentromeric region (~154 K_B_T/mm^2^) is significantly higher than those of loci at the interior (~110 K_B_T/mm^2^) and near-telomeric regions (~97 K_B_T/mm^2^) on chromosome 19 q arm. Similarly, we found the locus PR1 (pericentromeric region) on chromosome 19 p arm possess a high effective spring constant (~336 K_B_T/mm^2^) (Figure 3C and Table S3). We also calculated the effective diffusion constant *D_eff_* of loci (Supplementary Information, Figure 3D and Table S3) and found that the higher the effective spring constant of locus is, the lower its mobility is. Indeed, the effective external force constrains the locus dynamics. The effective spring constant of locus is expected to inversely correlate with locus territory. Polymer model predicts the relationship <*K_eff_*> = a <*R_g_^2^*>^-b^ with a=2 and b=1 (26). Our data showed that (a_PR1_, a_PR2_, a_LE_, a_LH_, a_LA_, a_T2_) = (2.09, 2.11, 2.69, 2.44, 2.65, 2.44) and (b_PR1_, b_PR2_, b_LE_, b_LH_, b_LA_, b_T2_) = (0.99, 0.99, 0.92, 0.95, 0.93, 0.94), consistent with the theoretical prediction (Figure 3E and Figure S5). To extract the information about inter-locus interactions and the tethering interaction of a locus to the local microenvironment, we modeled the chromosome by using the Rouse polymer model (27). For simplicity, we consider the polymer chain with one tethered locus at the position c_t_ and two labeled loci at the positions c_1_ and c_2_ (Figure 3F). In this model, *k_eff,i_* = *k_t_* K_R_ /(K_R_ +D_c_i*t*__*k_t_*) where K_R_ is the end-to-end effective spring constant of the polymer chain and D_c_i*t*__ is the distance between c_t_ and c_i_ (26). For the locus pair, LA/T2, we obtained the average end-to-end effective spring constant K_R_ ~ 25.0 (K_B_T/mm^2^) for chromosome 19 q arm and *k_t_* ~ 59.3 (K_B_T/mm^2^) where K_B_ is the Boltzmann constant and T is the absolute temperature of the environment (Supplementary Information). For the tethering spring constant near the pericentromeric region, we obtained *k_t_* ~ 193.5 (K_B_T/mm^2^) and 609.7 (K_B_T/mm^2^) at neighboring tethered loci of PR1 and PR2, respectively by assuming the same value of K_R_ and D_c_i*t*__ =1Mb. The simulation of b-polymer model (28) showed that the long-range interaction among loci contributes to the effective spring constant which is inversely proportional to D_c_i*t*__ (Figure 3 in (26)). Here, the genomic distance of the pair of loci LA/T2 is considerably small compared to the whole chromosome 19 so that the long-range interaction within a polymer chain can be neglected for the calculation of effective spring constants. Our results suggest that the tethering interactions of loci at the centromeric region are stronger than those of loci at the telomeric region.

**Figure 3.**
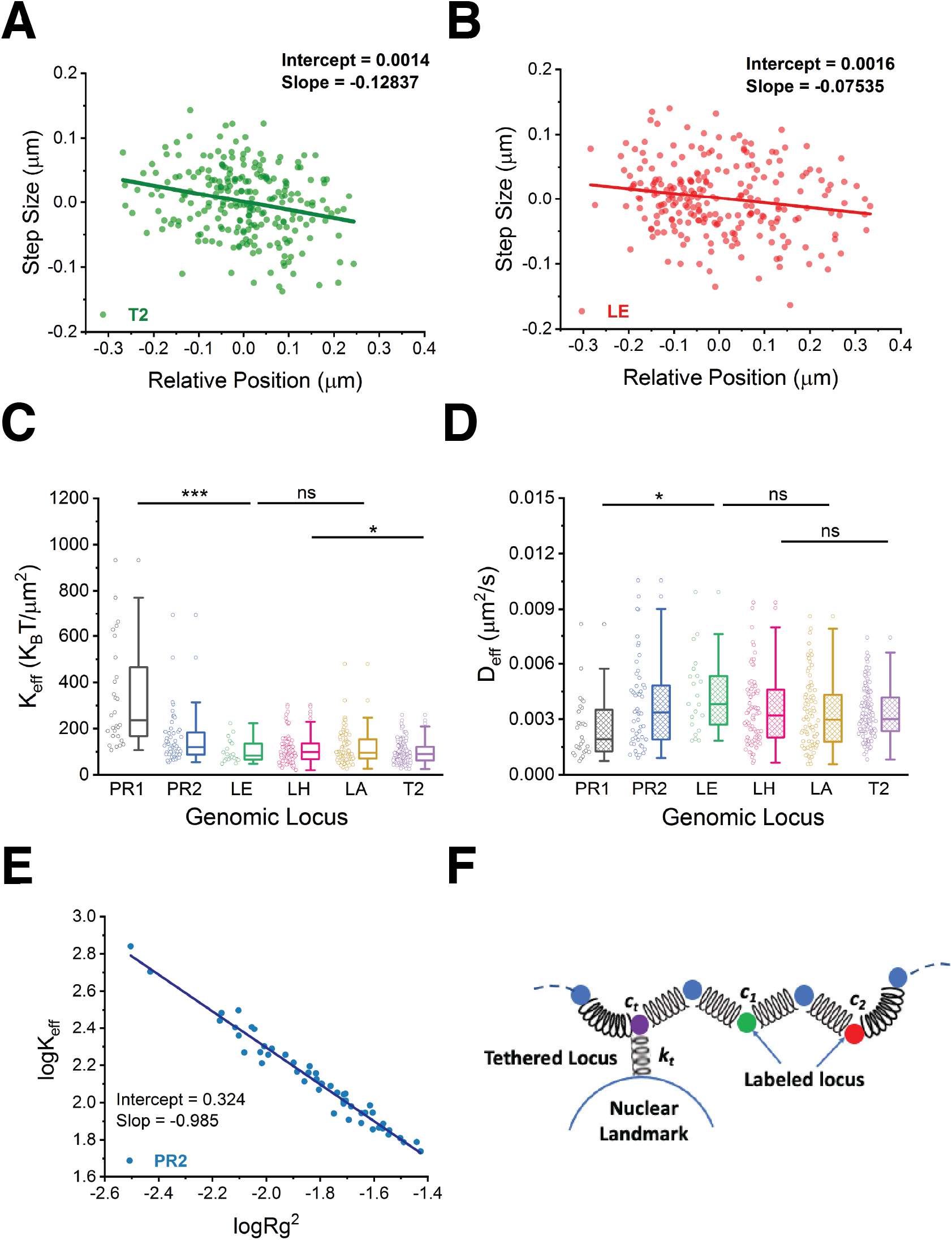
Dynamics and tethering of loci along human chromosome 19. (A)-(B) Scatter plots of step size versus the relative position to the centroid of locus trajectory for loci T2 and LE shown in **Figure 1.** (D). The data was fitted by linear regression (shown by the solid line). (C) Box-and-Whisker plot of effective spring constants (n = 28 trajectories for PR1, 52 trajectories for PR2, 21 trajectories for LE, 77 trajectories for LH, 77 trajectories for LA, and 127 trajectories for T2). (D) Box-and-Whisker plot of effective diffusion constants (numbers of trajectories are the same as in (C)). (E) A scatter plot of the effective spring constant versus the locus territory of locus PR2 in a log-log scale. The data was fitted with linear fit (the solid line) (F) Schematic of a Rouse polymer chain with a tethered locus ct and two labeled loci c_1_ and c_2_. The tethering spring constant is denoted by k_t_. Significance tests were performed using one-way ANOVA test: “ns” indicates not statistically significant (*p* > 0.05), “*” indicates significant difference *p* < 0.05, “**” indicates significant difference *p* < 0.005, and “***” indicates significant difference *p* < 0.0005.

### Conclusions and Perspectives

We have shown, by powerful CRISPR-Sirius live-cell real-time locus tracking, that the biophysical properties of genomic loci vary along a single chromosome (Figure 4), reflecting the inhomogeneity of chromatin folding and tethering to its surrounding microenvironment. We characterized chromatin compaction by the scaling (compaction) exponent of the power-lower relationship between the spatial distance and genomic distance of loci pairs. The compaction exponent decreased along with the length scale of chromatin and gradually become plateaued at the large genomic scale ~ 20 Mb (Figure 4B) consistent with the phenomenon observed on chromosome 19 p arm (20). This similarity might suggest the folding of chromosome p and q arms follows the same mechanism. However, we found, for the first time, the difference in the compaction level of chromosome 19 p arm and q arm in live human cells. The tighter folding of the q arm may result from its less number or fraction of active genes, compared to the p arm, shown by our RNA-seq analysis (Figure 2C). These results motivate us to hypothesize that both arms of a chromosome are folded by the same mechanism but regulated by its unique transcription profile. Further experiments will be required to test this hypothesis.

**Figure 4.**
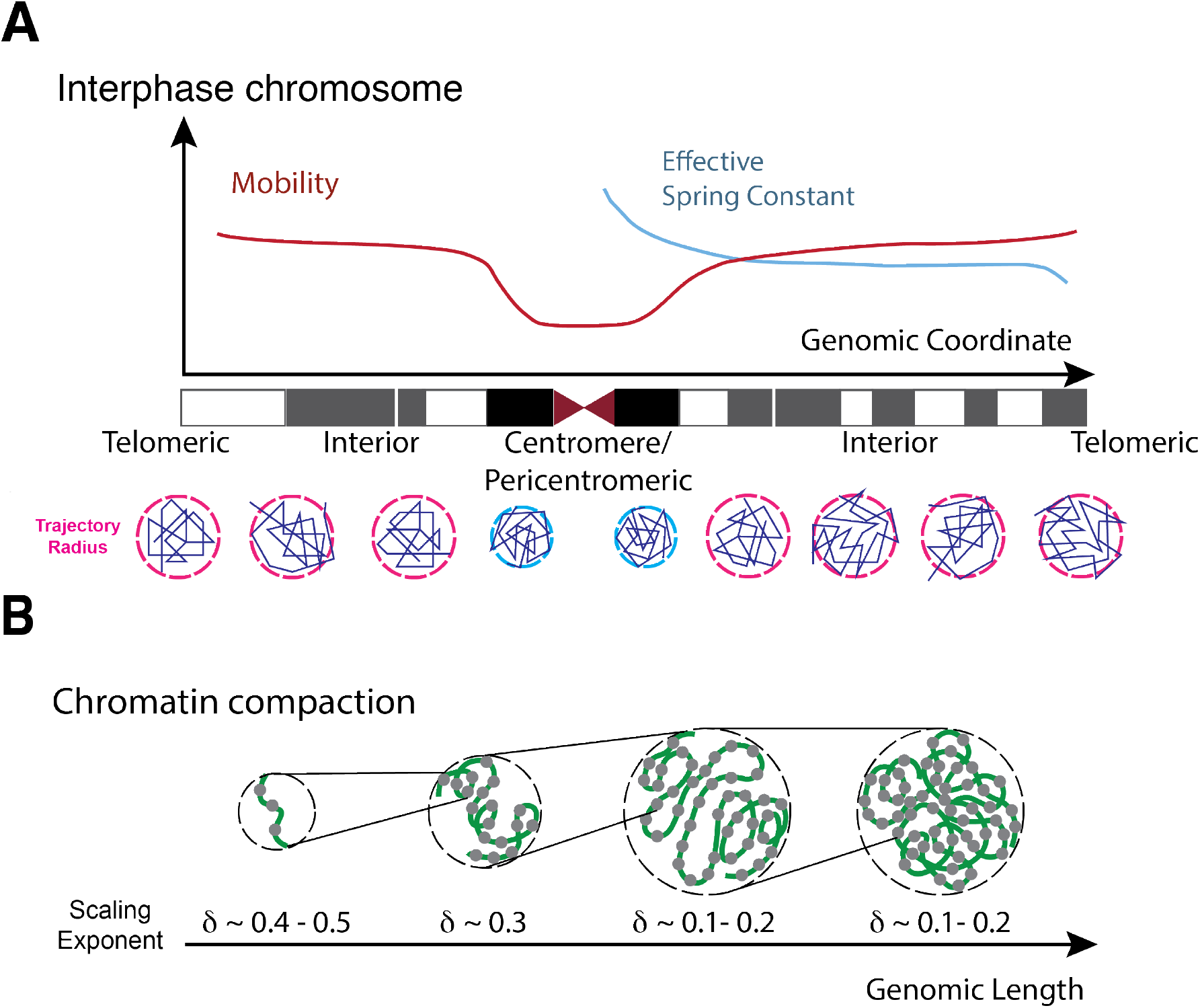
Summary diagram of single chromosome dynamics. (A) Variation of mobility and effective spring constants of genomic loci along the chromosome coordinate. (B) Compaction scaling exponents vary at different genomic length scale.

The mobility of loci, determined by MSD curve, also varies along the chromosome. We found that the dynamics of loci are subdiffusive with diffusion exponents 0.32 ~ 0.46 and the locus mobility increased from the centromeric region to the interior region and slightly went up at the telomeric region. Combining the low mobility of locus PR1 measured in this work with the previous results (20), a similar variation of locus mobility is also found on chromosome 19 p arm (Figure 4A). In addition, we found that the locus territory, defined as the area coved by the trajectory of locus movement, increases from the centromeric region to the interior region and becomes plateaued at the telomeric region on both arms of chromosome 19 (Figure 4B). To better understand how local forces constrain the locus dynamics, we modeled the short-time locus dynamics by the Langevin equation with an effective external force obeying Hooke’s law with an effective spring constant and applied on the locus. Using CRISPR-Sirius locus tracking data, we obtained the effective spring constant of locus and found that the effective spring constants vary along the chromosome. The loci at the pericentromeric region have the highest effective spring constant which then decays until reaches a plateau at the interior region and decreases again at the near-telomeric region. The variation of effective spring constants is observed to be inversely correlated to that of the effective diffusion constant of locus dynamics. By calculating the power-law decay of the effective spring constant versus locus territory and comparing it with polymer model prediction, we concluded that the locus dynamics is constrained by effective local force. To further decipher the effective force applied on loci, we modeled the chromatin elasticity and tethering interactions between loci and nuclear-landmarks by the Rouse model. The results show that the tethering interaction of loci at the centromeric region is stronger than that of loci at the telomeric region. We also observed that these loci with strong tethering couplings have high frequencies of association with nucleoli.

Using CRISPR-Sirius, we show that biophysical parameters characterizing chromatin dynamics and folding vary along a single chromosome. Contrary to averaged genome-wide chromatin dynamics in a whole nucleus, our measurement of locus dynamics at a single chromosome level can provide refined information about local interactions of loci and study the variety of chromatin dynamics among chromosomes. Combining with polymer models, CRISPR-Sirius opens a new window to study the interactions between chromatin and nuclear landmarks (e.g., nuclear bodies or nuclear lamina) and interactions between gene regulatory elements that reside on chromosomal DNA (e.g., enhancer-promoter interactions).

## Materials and Methods

### Plasmid Construction

The sgRNA sequences and genomic coordinates are listed in Table S4. sgRNAs were ordered from IDT or Sigma Oligo services and inserted into the sgRNA vectors using BbsI restriction sites. The expression vector, pHAGE-TO-dCas9-P2A-HSA, for dCas9 (nuclease-dead) from *S. pyogenes* was from our previous work in (16). PCP-GFP and pHAGE-EFS-MCP-HaloTag were previously described (6, 16). The expression vectors for dual guide RNAs, pPUR-P2A-BFP-mU6-sgRNA-Sirius-8XMS2 or pPUR-P2A-BFP-hU6-sgRNA-Sirius-8XPP7, were based on the pLKO.1 lentiviral expression system and described in (16). All the dCas9 and guide RNA expression vectors reported here are available on Addgene.

### Cell Culture, Lentivirus Production and Transduction

Human osteosarcoma U2OS cells were cultured on 35 mm glass-bottom dishes at 37°C in Dulbecco-modified Eagle’s Minimum Essential Medium (DMEM) containing high glucose and supplemented with 10% (vol/vol) fetal bovine serum. We used U2OS^dCas9-HSA/PCP-GFP/MCP-HaloTag^ cell line that was generated in our previous work for this study (20). Lentiviral particles that carry sgRNA plasmids were generated using HEK293T cells using the same protocol described in the reference (20). HEK293T cells were maintained in Iscove’s Modified Dulbecco’s Medium containing high glucose and supplemented with 1% GlutaMAX, 10% fetal bovine serum and 1% each penicillin and streptomycin. 24 hours before transfection, approximately 5×10^5^ cells were seeded in 6-well plates. For each well, 0.5 μg of pCMV-dR8.2 dvpr (Addgene), 0.3 μg of pCMV-VSV-G (Addgene), each constructed to carry HIV LTRs, and 1.5 μg of the plasmid containing the gene of interest were cotransfected by using TransIT transfection reagent (Mirus) according to manufacturer’s instructions. After 48 hours, the virus was collected by filtration through a 0.45 μm polyvinylidene fluoride filter. The virus was immediately used or stored at −80 °C. For lentiviral transduction, U2OS cells maintained as described above were transduced by Spinfection in 6-well plates with lentiviral supernatant for 48 hours and ~2×10^5^ cells were combined with 1 ml lentiviral supernatant and centrifuged for 30 minutes at 1200 x *g*.

### Analysis of RNA-seq Data

The raw reads of the RNA-seq for osteosarcoma (U2OS) cells were obtained from Sequence Research Archive (GEO accession no. GSE118488, SRA - SRX4549306 and SRX4549307). The quality of the raw data was assessed using the FastQC v. 0.72 (29). Raw data was aligned to the human genome build GRCh38 using the bioinformatic tool HISAT2 v. 2.1.0 (30), which generated the Binary Alignment Map (BAM). The reads in the generated (BAM) were then counted using FeatureCounts v.1.6.4 (31). The raw read counts and read length corresponding to each gene were used to compute TPM (Transcripts Per Million) values (32). To identify the active genes, we used the DAFS algorithm (33) based on Kolmogorov Smirnov distance statistics (34) to calculate the TPM cutoff for active genes in the RNA seq data set. The DAFS algorithm used is based on model-based clustering and uses the R package mclust (35) and earth (36) to predict the cut-off value corresponding to active genes. After identifying active genes, we used R package TxDb.Hsapiens.UCSC.hg38.knownGene to obtain the genomic coordinates corresponding to each active gene on chromosome 19(37). The genomic coordinates were finally used to determine the number of genes on the short arm (p arm) and long arm (q arm) of chromosome 19.

### Fluorescence Microscopy

An Olympus IX83 microscope was equipped with three EMCCD cameras (Andor iXon 897), four lasers (405 nm, 488 nm, 561 nm, and 647nm), mounted with a 1.6x magnification adapter and 60x apochromatic oil objective lens (NA 1.5, coverslip- and temperature-corrected), resulting in a total of 96x magnification. The microscope stage incubation chamber was maintained at 37 °C with CO2 and humidity supplement. A laser quad-band filter set for TIRF (emission filters at 445/58, 525/50, 595/44, 706/95) was used to collect fluorescence signals simultaneously. Imaging data were acquired by CellSens software. The localization precision was measured by capturing 120 frames of 0.1 μm coverglass-immobilized TetraSpeck fluorescent microspheres (n = 220 for 16 seconds and n = 218 for 80 seconds) at 100 ms exposure time and calculating the standard deviations, a method developed by Jeff Gelles (38). As described in the main text, the localization precision was ~5 nm in 4 seconds, ~ 6 nm in 16 seconds, and ~ 10 nm in 80 seconds (Figure S1 and S2) Image size was adjusted to show individual nuclei and intensity thresholds were set on the basis of the ratios between nuclear foci signals to background nucleoplasmic fluorescence. To quantify the spatial distance or track the dynamics, only pairs of loci lying in the same focal plane were analyzed.

### Imaging Processing

The images were registered and analyzed by *Fiji* (39) and *Mathematica* (Wolfram) software. Images from the green and red channels were registered by 0.1 μm coverglass-absorbed TetraSpeck fluorescent microspheres (Invitrogen) as a standard sample. To eliminate the movement from live cells, localizations of individual genomic loci were calibrated by the motion relative to the nuclear centroid. Detailed calculation of the mean-square displacement (MSD), relative spatial distance (D), gyration radius of trajectory radius (R_g_), effective spring constant (K_**eff**_), tethering coupling and position analysis are described in (20) and Supplementary Information. All analyses were performed by *Mathematica* and graphs were generated by *OriginPro* (OriginLab version 2019b). All box plots were generated using the default setting of the *OriginPro*. Box spans from first to last quartiles and whisker length is determined by the outermost data point that falls within the upper inner and lower inner fence (a coefficient =1.5). Significance tests were performed using one-way ANOVA function in OriginPro. “ns” denotes for not statistically significant (*P* > 0.05), * indicates *P* < 0.05, ** indicates *P* < 0.005, and *** indicates *P* < 0.0005.

## Data Availability

The RNA-seq data is an available resource on Sequence Research Archive (GEO accession no. GSE118488, SRA –SRX4549306 and SRX4549307) deposited by Rachel Litman Flynn (40). All other data presented in this paper are available upon reasonable request.

## Acknowledgements

We thank Sydney Willey and Riley Moran in the Tu lab for the help in part of plasmid construction. This work was supported by the NIH grant R00 GM126810 and OSU start-up fund to L-C.T. We are also very grateful to Mark Parthun and Daniel Schoenberg at OSU Medical School for sharing lab equipment and strong encouragement during the study.

## Author Contributions

L.-C. Tu, and Y.-C. Chung conceived the project and designed the experiments; Y.-C. Chung conceived and performed the mathematical treatment and calculation of the dynamics results; M. Bisht performed RNA seq analysis; Y.-C. Chung and L.-C. Tu performed experiments; Y.-C. Chung and L.-C. Tu performed imaging processing and quantification analysis; L.-C. Tu, Y.-C. Chung, interpreted data and wrote the paper.

## Supplemental Information

Supplemental Information includes five figures, four tables, and one movie.

---

**Table S1.** Biophysical parameters extracted from single locus trajectories.

| Genomic Locus | MSD Exponent ($\beta$) | Radius ($\mu\text{m}$) | $D_{app}$ ($\mu\text{m}/\text{sec}^\beta$) | N (Trajectory) |
| --- | --- | --- | --- | --- |
| T2 | $(4.57 \pm 0.02) \times 10^{-1}$ | $(1.52 \pm 0.37) \times 10^{-1}$ | $(3.02 \pm 0.10) \times 10^{-3}$ | 127 |
| LA | $(4.04 \pm 0.03) \times 10^{-1}$ | $(1.46 \pm 0.42) \times 10^{-1}$ | $(3.36 \pm 0.16) \times 10^{-3}$ | 77 |
| LH | $(4.17 \pm 0.02) \times 10^{-1}$ | $(1.51 \pm 0.46) \times 10^{-1}$ | $(3.35 \pm 0.11) \times 10^{-3}$ | 77 |
| LE | $(3.27 \pm 0.04) \times 10^{-1}$ | $(1.51 \pm 0.32) \times 10^{-1}$ | $(4.43 \pm 0.23) \times 10^{-3}$ | 21 |
| PR2 | $(3.59 \pm 0.01) \times 10^{-1}$ | $(1.28 \pm 0.32) \times 10^{-1}$ | $(2.98 \pm 0.05) \times 10^{-3}$ | 52 |
| PR1 | $(3.57 \pm 0.07) \times 10^{-1}$ | $(0.88 \pm 0.26) \times 10^{-1}$ | $(1.47 \pm 0.15) \times 10^{-3}$ | 28 |
| IDR1* | $(3.41 \pm 0.08) \times 10^{-1}$ | $(1.33 \pm 0.51) \times 10^{-1}$ | $(5.31 \pm 0.12) \times 10^{-3}$ | 7 |
| IDR2* | $(3.61 \pm 0.08) \times 10^{-1}$ | $(1.52 \pm 0.33) \times 10^{-1}$ | $(4.19 \pm 0.10) \times 10^{-3}$ | 19 |
| IDR3* | $(3.64 \pm 0.03) \times 10^{-1}$ | $(1.48 \pm 0.36) \times 10^{-1}$ | $(4.86 \pm 0.05) \times 10^{-3}$ | 49 |
| TCF3* | $(4.60 \pm 0.05) \times 10^{-1}$ | $(1.28 \pm 0.44) \times 10^{-1}$ | $(2.88 \pm 0.04) \times 10^{-3}$ | 12 |
| IDR4* | $(4.41 \pm 0.07) \times 10^{-1}$ | $(1.58 \pm 0.52) \times 10^{-1}$ | $(4.16 \pm 0.08) \times 10^{-3}$ | 11 |

\* These data were adopted from (5).

**Movie S1.** This movie shows a typical movement of the locus pair LH/T2 in the U2OS cell nucleus recorded over 80 seconds (120 image frames). LH is labeled in red by Halo tag-JF549 and T2 is labeled by GFP. Movie play rate is 30 Hz. The trajectories of loci in this movie are shown in Figure 1E.

